# A Bayesian Monte Carlo approach for predicting the spread of infectious diseases

**DOI:** 10.1101/617795

**Authors:** Olivera Stojanović, Johannes Leugering, Gordon Pipa, Stéphane Ghozzi, Alexander Ullrich

**Author notes:** These authors contributed equally to this work (Olivera Stojanović, Johannes Leugering). Corresponding authors (OS), (AU).

## Abstract

In this paper, a simple yet interpretable, probabilistic model is proposed for the prediction of reported case counts of infectious diseases. A spatio-temporal kernel is derived from training data to capture the typical interaction effects of reported infections across time and space, which provides insight into the dynamics of the spread of infectious diseases. Testing the model on a one-week-ahead prediction task for campylobacteriosis and rotavirus infections across Germany, as well as Lyme borreliosis across the federal state of Bavaria, shows that the proposed model performs on-par with the state-of-the-art *hhh4* model. However, it provides a full posterior distribution over parameters in addition to model predictions, which aides in the assessment of the model. The employed Bayesian Monte Carlo regression framework is easily extensible and allows for incorporating prior domain knowledge, which makes it suitable for use on limited, yet complex datasets as often encountered in epidemiology.

**Author summary:** **Why was this study done?**

- Statistical modeling is invaluable to public-health policy as it helps understand and anticipate the dynamics of the spread of infectious diseases. The available training data is often limited and reported with a low spatial and temporal resolution. This poses a challenge and makes it particularly important to incorporate domain knowledge and prior assumptions to guide the modeling process.
- In order to evaluate the trustworthiness and reliability of a model’s predictions, it is crucial to be able to interpret the model and quantify the model uncertainty.
- To address this, we develop an interpretable model that uses Bayesian inference (rather than commonly used maximum likelihood estimation) and provides a probability distribution over inferred parameters.

**What did the researchers do and find?**

- We develop and test a single probabilistic model that learns to predict the number of weekly case counts for three different diseases (campylobacteriosis, rotaviral enteritis and Lyme borreliosis) at the county level one week ahead of time.
- We employ a Bayesian Monte Carlo regression approach that provides an estimate of the full probability distribution over inferred parameters as well as model predictions.
- The model learns an interpretable spatio-temporal kernel that captures typical interactions between infection cases of the tested diseases.
- The predictive performance of our model compares favorably with a contemporary reference model for all diseases tested.

**What do these findings mean?**

- Interpretable predictive models can be applied to surveillance data to gain insights into the dynamics of infectious diseases.
- Probabilistic modeling approaches provide a suitable framework for many challenges of working with epidemiological data.

## Introduction

Public-health agencies have the responsibility to *detect, prevent* and *control* infections in the population. In Germany, the Robert Koch Institute collects a wide range of factors, such as location, age, gender, pathogen, and further specifics, of laboratory confirmed cases for approximately 80 infectious diseases through a mandatory surveillance system [1]. Since 2015, an automated outbreak detection system, using an established aberration detection algorithm [2], has been set in place to help *detect* outbreaks [3, 4]. However, *prevention* and *control* require quantitative *prediction* instead of mere *detection* of anomalies and thus prove more challenging. For logistical, computational and privacy reasons, epidemiological data is typically reported or provided in bulk, often grouped by calendar weeks and counties. Predictions thus have to be made about the number of cases per time-interval and region, based on a history of such measurements.

Since outbreaks can extend over multiple counties, states or even nations, spatio-temporal models are typically employed. Some approaches use scan statistics to identify anomalous spatial or spatio-temporal clusters [5, 6], while others model and predict case counts as time series or point processes [7]. A major advantage of such predictive models over univariate aberration detection approaches is the additional insight they can provide into the factors contributing to the spread of infectious diseases.

In general, we distinguish four qualitatively different classes of predictive features: *spatial, temporal, spatio-temporal* and *(spatio-temporal) interaction* effects. The former three are purely functions of space, time or both, modeling *seasonal* fluctuations and *trends, geographical* influences or localized time-varying effects, such as *region-specific demographics* or *legislation*, respectively. The latter is an autoregressive variable that captures how an observed infection influences the number of further infections in its neighborhood over time, which depends on differences in *patients’ behavior, transmission vectors, incubation times* and *duration* of the respective diseases. Even in the absence of direct contagion, previously reported cases can provide valuable *indirect* information for predicting future cases through latent variables. The effect on the expected number of cases at a given place and time due to interactions can thus be expressed as a (unknown) function of spatial and temporal distance to previously reported cases. Particularly for regions with less available historic data or those strongly influenced by their neighbours, e.g. smaller counties close to larger cities [8], incorporating the county’s and its neighbours’ recent history of case counts can improve predictions.

The state-of-the-art spatio-temporal *hhh4* method [7] [9] assumes aggregated case counts to follow a Poisson or Negative Binomial distribution around a mean value determined by “epidemic” and “endemic” components. The epidemic component can capture the influence of previous cases from the same or neighbouring counties, e.g. potentially weighted by the counties’ adjacency order, while the endemic component models the expected baseline rate of cases.

For *not aggregated* data, the more general *twinstim* method [7] models the interaction effects due to individual cases by a self-exciting point process with predefined continuous spatio-temporal kernel, rather than through a binary neighborhood relation as in the *hhh4* model. Optimizing such a kernel for a specific dataset provides an opportunity to incorporate or even infer information about the infectious spread of the disease at hand.

In the following, we present a Bayesian spatio-temporal interaction model (referred to as BSTIM), as a synthesis of both approaches: a probabilistic generalized linear model (GLM) [10] predicts aggregated case counts within spatial regions (counties) and time intervals (calendar weeks) using a history of reported cases, temporal features (seasonality and trend) and region-specific as well as demographic information. Like for the *twinstim* method, interaction effects are modeled by a continuous spatio-temporal kernel, albeit parameterized with parameters inferred from data. Since the aggregated reporting of case counts per calendar week and county leaves residual uncertainty about the precise time and location of an individual case, we model times within the respective week and locations within the respective county as latent random variables. Monte Carlo methods are employed to evaluate posterior distributions of parameters as well as predictions, which are subsequently used to assess the quality of the model.

For three different infectious diseases, *campylobacteriosis, rotaviral enteritis* and *Lyme borreliosis*, the interpretability of the inferred components, specifically the interaction effect kernel, is discussed and the predictive performance is evaluated and compared to the *hhh4* method.

## Materials and methods

We evaluate both the proposed BSTIM as well as the *hhh4* reference model on a one-week-ahead prediction task, where the number of cases in each county is to be predicted for a specific week, given the previous history of cases in the respective as well as surrounding counties. Instead of point estimates, we are interested in a full posterior probability distribution over possible case counts for each county and calendar week – capturing both aleatoric uncertainty due to the stochastic nature of epidemic diseases as well as epistemic uncertainty due to limited available training data. The data for this study is provided by the Robert Koch Institute, and consists of weekly reports of case counts for three diseases, campylobacteriosis, rotavirus infections and Lyme borreliosis. They are aggregated by county^1^ and collected over a time period spanning from the 1st of January 2011 (2013 for borreliosis) to the 31st of December 2017 via the *SurvNet* surveillance system [1]. Aggregated case counts of diseases with mandatory reporting in Germany can be downloaded from https://survstat.rki.de. For each of the three diseases, the data preceding 2016 is used for training the model, while the remaining two years are used for testing. A software implementation of the BSTI Model presented here is available online at https://github.com/ostojanovic/BSTIM.

### The BSTI Model

The proposed model is optimized to predict the number of reported cases in the future (e.g. the next week), based on prior case counts. For modeling purposes we assume counts are distributed as a Negative Binomial random variable around an expected value *µ*(*t, x*) that varies with time (*t*) and space (*x*). We further assume that the relationship between each feature *f*_*i*_(*t, x*) and the expected value *µ*(*t, x*) can be expressed in a generalized linear model of the Negative Binomial random variable *Y* (*t, x*) using the canonical logarithmic link function. For the limited available data, an appropriate choice of priors is crucial to prevent overfitting. We use half-Gaussian priors for interaction effects to regularize the coefficients while ensuring positivity of the inferred kernel. A half-Cauchy distribution is used as a weakly informative prior [11] to enforce positivity of the dispersion parameter of the residual Negative Binomial distribution. During training, the model parameters are regularized by a Gaussian prior for the weights of all features. Since both the basis functions (c.f. section *Interaction effects*) and the coefficients used for modeling the interaction effects are nonnegative, the resulting interaction kernel is thus also constrained to be nonnegative.

The full probabilistic model for training can thus be summarized as follows:

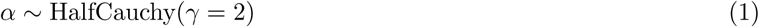

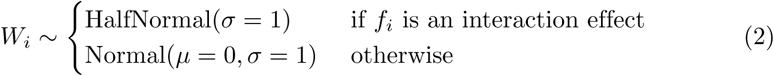

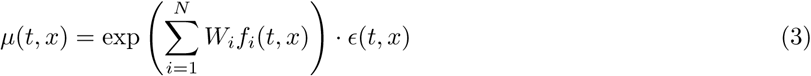

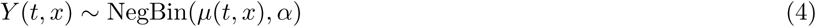

where:

*α* is a dispersion parameter

*N* is the total number of used features

*W*_*i*_ are model weights

*f*_*i*_(*t, x*) are features varying in time and space

*ϵ*(*t, x*) is the exposure varying in time and space

*t* refers to a time-interval (i.e. one calendar week)

*x* refers to a spatial region (i.e. one county)

For prediction, the priors over the dispersion parameter and weights are replaced by the corresponding posterior distribution inferred on the training set.

A schema of our model is shown in Fig.1. To capture the interaction effects between different places over time, a continuous spatio-temporal kernel is estimated through a linear combination of 16 basis kernels. The individual contribution due to each of these basis kernels is included into the model as a feature. Four temporal periodic *basis functions* are used to capture seasonality and five sigmoid *basis functions* (one for each year of available training data) to capture temporal trends. Four region-specific features (ratio of population in a county belonging to three age groups and one political component) are used, which results in 29 features. In addition, the logarithm of the population of each county in the respective year is used as a scaling parameter (exposure) *ϵ*.

**Fig 1.**
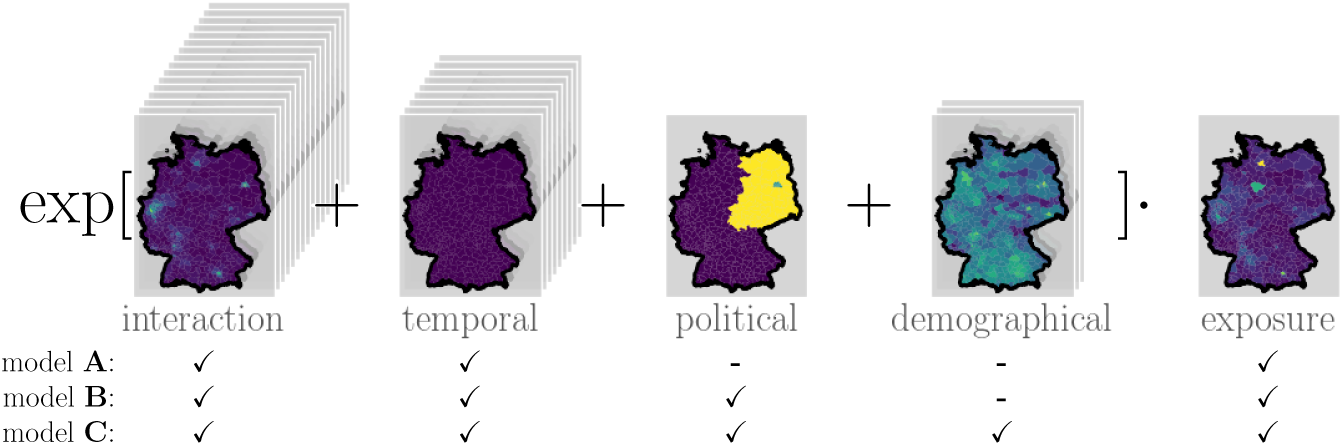
Model scheme. Exemplary contributions from different features, grouped into interaction, temporal, political and demographical components, each evaluated in all counties in Germany for campylobacteriosis in the week 30 of 2016. Each county’s total population is always included as an exposure coefficient. We consider three models of increasing complexity, A, B and C, that differ in whether features are included (✓) or not (-).

For example, given one parameter sample *w* = [*w*_1_, …, *w*_*n*_], inferred from the training set of campylobacteriosis case counts, the conditional mean prediction within county *x* during week *t* is determined as follows:

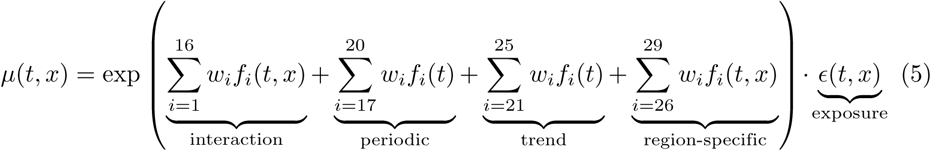

### Monte Carlo sampling procedure

The model described above determines the posterior distribution over parameters by the data-dependent likelihood and the choice of priors. We want to capture this parameter distribution in a fully Bayesian manner, rather than summarize it by its moments (ie. mean, covariance, etc.) or other statistics. Since an analytic solution is intractable, we use Markov Chain Monte Carlo (MCMC) methods to generate unbiased samples of this posterior distribution. These samples can be used for evaluation of performance measures (here Widely Applicable Information Criterion (WAIC), and Dawid-Sebastiani score; cf. section *Predictive performance evaluation and model selection*), visualization or as input for a superordinate probabilistic model.

Our model combines features that can be directly observed (e.g. demographic information) with features that can only be estimated (e.g. interaction effects, due to uncertainty caused by data aggregation). To integrate the latter into the model, we generate samples from the distribution of interaction effects features as outlined in section *Interaction effects*.

The sampling procedure generates samples from the *prior* distribution over parameters and combines them with training data and our previously generated samples of the interaction effect features to produce samples of the *posterior* parameter distribution. These samples from the inferred joint distribution over *parameters* are then used to generate samples of the posterior distribution of model *predictions* for testing data.

We employ a Hamiltonian Monte Carlo method, No-U-Turn-Sampling [12], implemented in the probabilistic programming package *pyMC3* [13].

#### Interaction effects

Each reported case provides valuable information about the expected number of cases to come in the near future and close proximity. We suppose that this effect of an individual reported infection on the rate of future (reported) infections in the direct neighborhood can be captured by some unknown function *κ*(*d*_time_(*t*_⋆_, *t*_*k*_), *d*_geo_(*x*_⋆_, *x*_*k*_)), which we refer to as *interaction effect kernel* in the following, where (*t*_*k*_, *x*_*k*_) refer to the time and location of the *k*-th reported case and (*t*_⋆_, *x*_⋆_) represent the time and location of a hypothetical future case. Here, *d*_geo_(*x, y*) represents the geographical distance between two locations *x* and *y*, whereas *d*_time_(*t, s*) denotes the time difference between two time points *t* and *s*. Thus, *κ*(·, ·) is a radial, time- and location-invariant kernel, depending only on the spatial and temporal proximity of the two (hypothetical) cases. For the sake of simplicity, we assume that interaction effects due to individual infections add up linearly.

Since *κ* is not known a-priori for each disease, we wish to infer it from data. To this end, we approximate it by a linear combination of spatio-temporal basis kernels *κ*_*i,j*_ with coefficients *w*_*i,j*_ that can be inferred from training data:

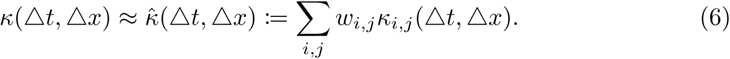

As the basis functions for the interaction effect kernel, we choose the products 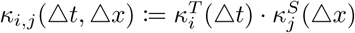 between one temporal 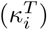 and one spatial factor 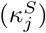, each (cf. Fig.2). As temporal factors, we use the second order B-spline basis functions corresponding to the knot vector [0, 0, 1, 2, 3, 4, 5] (measured in weeks), resulting in four smooth, positive, unimodal functions, spanning the overlapping time interval from zero to two weeks, zero to three weeks, one to four weeks and two to five weeks after a reported case, respectively. Outside these intervals, the functions are identically zero; a-causal effects, i.e. the influence of a reported case on hypothetical other cases reported at an earlier time, are thus excluded. As spatial factors, we use exponentiated quadratic kernels (i.e. univariate Gaussian functions) centered at a distance of 0km to a reported case, with shape parameters *σ* of 6.25km, 12.5km, 25.0km, and 50.0km. Since both temporal as well as spatial factors are nonnegative, the resulting basis functions *κ*_*i,j*_ are also nonnegative, and, consequently, a linear combination with nonnegative weights *w*_*i,j*_, as enforced in the fitting procedure, must result in a nonnegative interaction effect kernel 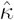. See Fig.2 for an illustration of how the basis functions *κ*_*i,j*_ are constructed.

**Fig 2.**
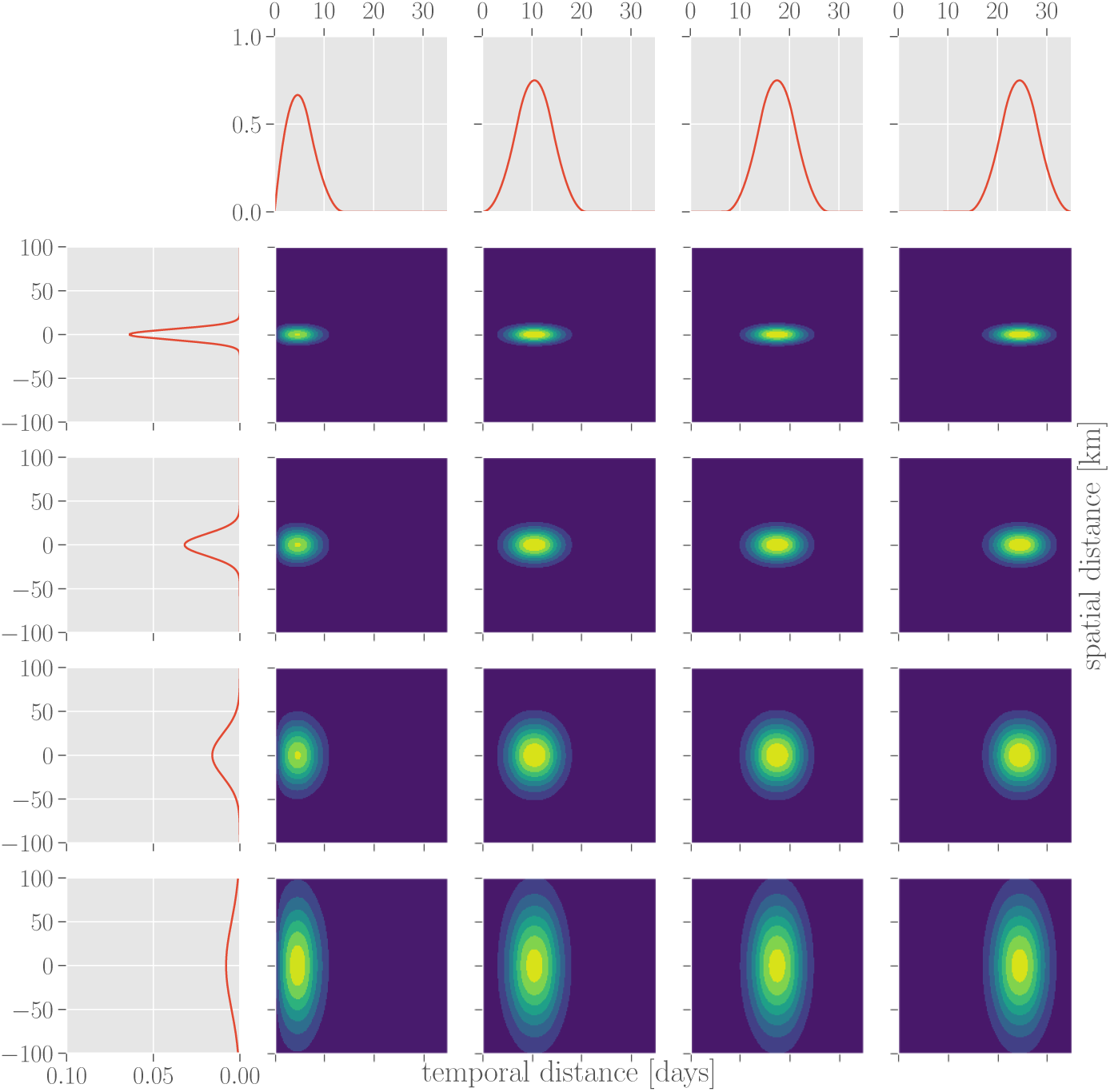
Spatial and temporal basis functions for interaction kernel. The inferred interaction kernel is composed of a linear combination of spatio-temporal basis functions (four-by-four grid of contour plots), each of which is a product of one spatial (left column) and one temporal factor (top row).

Since the contributions of individual cases are assumed to sum up linearly, the total influence of all cases that were previously reported at times and places (*t*_*k*_, *x*_*k*_), *K* ∈ 1 …*n* onto the expected rate of cases reported at a later time *t* and location *x* is given by:

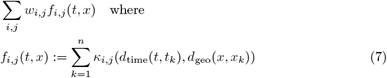

Each *f*_*i,j*_(*t, x*) for *i ∈* 1, …, 4, *j ∈* 1, …, 4 is a spatio-temporal function that depends on all cases reported prior to *t*, providing us with a total of 16 autoregressive features to use for the model. By determining the corresponding coefficients *w*_*i,j*_, the fitting procedure thus allows us to infer an interaction effect kernel 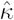 in a 16-dimensional parameterized family from data. It should be noted, however, that since the basis functions *κ*_*i,j*_ capture strongly correlated and possibly redundant information, the effective number of degrees of freedom (as estimated by WAIC during model selection) may be well below 16. Since we work with aggregated data at a spatial resolution of counties and a temporal resolution of calendar weeks, the exact time and location of an individual case report, as well as time and location of a hypothetical future case, are conditionally independent random variables given the county and week in which they occur. Because of this epistemic uncertainty, the features *f*_*i,j*_ derived in equation 7 are thus random variables themselves. To deal with this uncertainty, the *twinstim* model proposed in [7] suggests to replace these features by their expected values, which can be numerically approximated efficiently. Here, instead of using such point-estimates, which might lead the model to underestimate its uncertainty, we want to incorporate the features *f*_*i,j*_ directly into our probabilistic model and thus need to account for their full probability distribution.

While this distribution is intractable to calculate analytically, we can generate unbiased samples from it through rejection sampling: For a case reported in a given calendar week and county, possible sample points of a precise time and location can be independently generated by uniformly drawing times from within the corresponding week and locations from a rectangle containing the county, rejecting points that fall outside the county’s boundary. By randomly drawing a sample time and location for each reported case, we can thus generate an unbiased sample of the (unavailable) data prior to aggregation that accurately reflects the uncertainty caused by the aggregation procedure. Using these resulting sample times and locations in place of *t*_*k*_ and *x*_*k*_ in equation 7 yields unbiased samples of the features *f*_*i,j*_, which are in turn used when generating samples of the model’s posterior parameter distribution (cf. section *Monte Carlo sampling procedure*).

It bears repeating that what we refer to as interaction effect features in this paper are thus in fact latent random variables due to the epistemic uncertainty caused by aggregated reporting of infections by counties and calendar weeks.

### Additional features

Infection rates vary in time due to natural processes, such as seasons and climate trends, evolution of pathogens and immunization of the population, as well as societal developments such as scientific and technological advancement and medical education. Within Germany these effects may not differ much across space and can thus be included into the model as feature functions *f*_*i*_ (*t*) that only depend on time. For modeling yearly seasonality, four sinusoidal basis functions (ie. sin (2*π · t · ω*_yearly_), sin (4*π · t · ω*_yearly_), cos (2*π · t · ω*_yearly_), cos (4*π · t · ω*_yearly_)) are used as temporal periodic components, where *ω*_yearly_ = (1 year)^-1^. Slower time-varying effects are subsumed in a general trend modeled by a linear combination of one logistic function (ie. 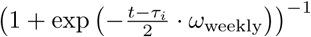) centered at the beginning of each year (*τ*_*i*_) with slope ½ *ω*_weekly_, where *ω*_weekly_ = (1 week)^-1^.

Due to the historical division between eastern and western Germany, and their different developments, some structural differences remain, such as unemployment rate, density of hospitals and doctors, population density, age structure etc. [14, 15] To account for such systematic differences, a political component, which we refer to as the *east/west component* in the following, is introduced which labels all counties that were part of the former German Democratic Republic as 1 and counties that were part of the Federal Republic of Germany as 0. Since Berlin itself was split into two parts, yet todays counties don’t accurately reflect this historic division, counties within Berlin are labeled with an intermediate value of 0.5.

Since diseases can affect children and elderly in different ways, yearly demographic information about each county is incorporated into the model. The logarithm of the fraction of population belonging to three age groups (ages [0 *-* 5), [5 *-* 20) and [20 *-* 65)) is used. The total population of each county acts as a scaling factor for the predicted number of infections^2^.

### Predictive performance evaluation and model selection

To evaluate the predictive performance of the model, forecasts of the number of infections are made one calendar week ahead of time for each disease and each county. To determine the relevance of different features, model selection is performed on the training dataset between three models of different complexity [Fig.1]:

**model A** - includes interaction and temporal (periodic and trend) components,

**model B** - includes interaction, temporal and political components,

**model C** - includes interaction, temporal, political and demographic components.

The Widely Applicable Information Criterion [16] is applied to the posterior distribution over parameters and predictions from the training set to determine which combination of features (i.e. model A, B or C) minimizes the generalization error while penalizing a large *effective* number of parameters. This is relevant here since modeling interaction effects introduces multiple features that capture redundant information.

Different error measures are applied to evaluate the fit of the predictive distribution for the test set to observations. Deviance of the Negative Binomial distribution (i.e. the expected difference between the log-likelihood of observations and the log-likelihood of the predicted means) is used as a likelihood-based measure and the Dawid-Sebastiani score (a covariance-corrected variant of squared error, cf. [17]) is included as a distribution-agnostic proper scoring rule.

To evaluate the performance of the model presented here as well as an *hhh4* model implementation for reference, we compare the resulting distributions of scores across counties.

### The *hhh4* model reference implementation

We use an *hhh4* model for Negative Binomial random variables, implemented in the R package “surveillance” [18], with a mean prediction composed of an epidemic and an endemic component. The epidemic component is a combination of an autoregressive effect (models reproduction of the disease within a certain region) and a neighborhood effect (models transmission from other regions). The endemic component models a baseline rate of cases due to the same features as described above. The reference model is trained and evaluated on the same datasets as the BSTIM.

## Results and discussion

Testing models of varying complexity (see Fig.1) reveals that the most complex model (model complexity C, including interaction effects, temporal, political as well as demographical features) generalizes best as measured by WAIC for all three different tested diseases (campylobacteriosis, rotavirus and borreliosis). [Tab.1] For the remainder of this text, we thus focus only on the full model variety C. The posterior parameter distribution inferred from the training data can be analyzed in itself, which provides valuable information about the disease at hand as well as the suitability of the model. Subsequently, it is used to generate one-week-ahead predictions for the test data.

**Table 1.** Training set WAIC scores for the three tested diseases and the three levels of model complexity.

| model | campylobacteriosis | rotavirus | borreliosis |
| --- | --- | --- | --- |
| **A** | 423422 | 349929 | 31499.1 |
| **B** | 420294 | 339715 | 31501.1 |
| **C** | 420151 | 338735 | 30810 |

### The inferred model

The procedure outlined above produces samples from the posterior parameter distribution, which in turn provides a probability distribution over interaction kernels. Due to the large number of free parameters (16) involved (see Fig.2), the family of parameterized kernels is flexible enough to capture different disease-specific interactions in time and space. The mean interaction kernel for campylobacteriosis (see Fig.3, 1A) shows the furthest spatial influence over up to 75 km, whereas rotavirus (see Fig.3, 2A) and borreliosis (see Fig.3, 3A) are more localized within a radius of up to 25 km. Borreliosis exhibits longer lasting interaction effects, extending up to four weeks. Despite the fact that borreliosis is not contagious between humans, this is consistent with a pseudointeraction effect due to a localized, slowly changing latent variable such as the prevalence of infected ticks or other seasonal factors.

**Fig 3.**
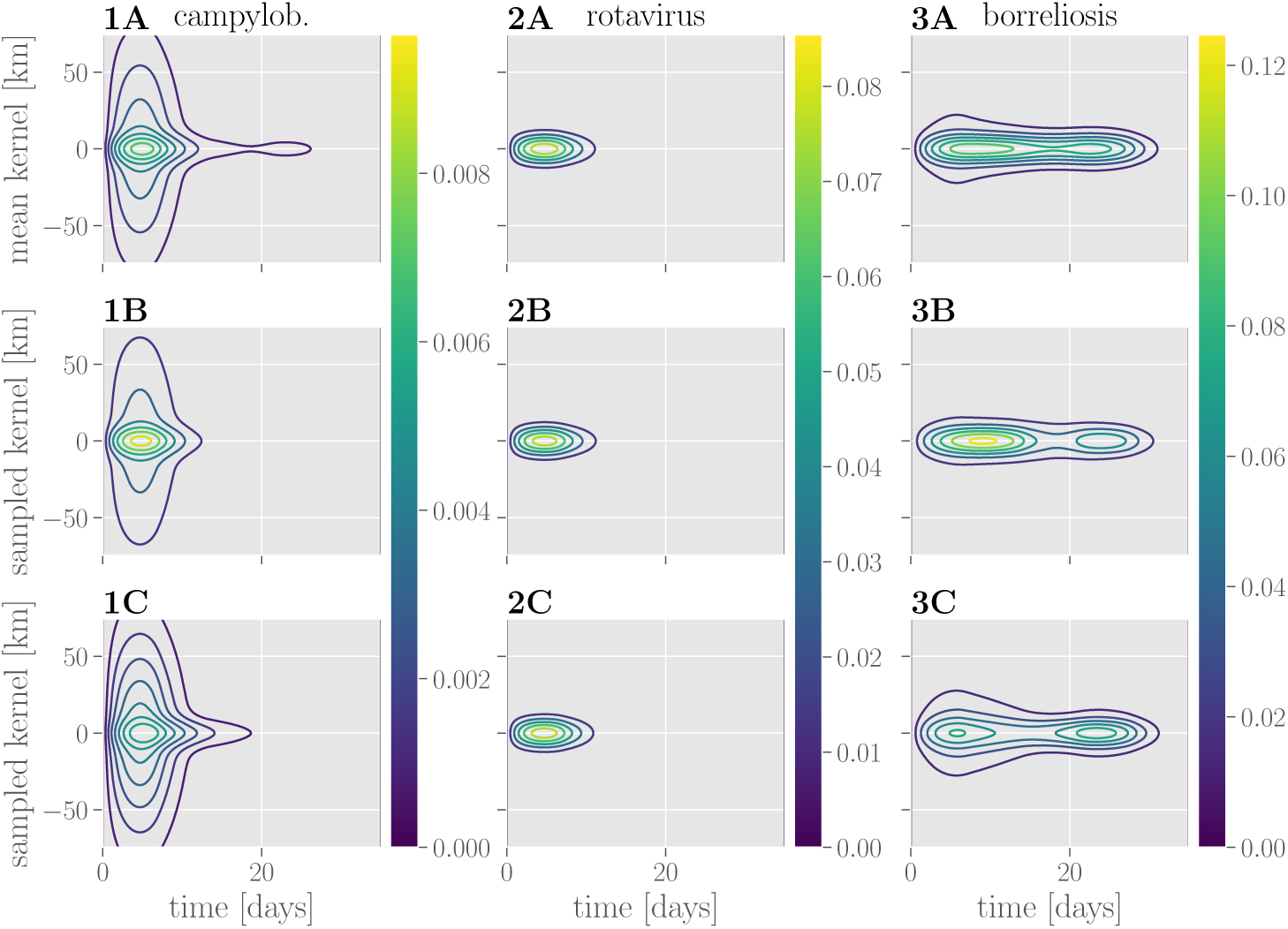
Learned interaction effect kernels. Kernels for campylobacteriosis are shown in **1A-C**, for rotavirus in **2A-C** and for borreliosis in **3A-C**. Mean interaction kernels are shown in the **row A**, while **rows B and C** show two random samples from the inferred posterior distribution over interaction kernels.

Looking at individual samples from the respective kernel distributions (see Fig.3, rows B and C) reveals a degree of uncertainty over the precise kernel shape for the different diseases: while there is little variation in the kernel shape inferred for rotavirus, there is uncertainty about the temporal profile of interactions for campylobacteriosis. See also supplementary figures S1 Fig, S2 Fig and S3 Fig for an overview of the respective posterior distributions over interaction effect coefficients.

The seasonal components (see Fig 4) for campylobacteriosis and borreliosis show a yearly peak in July and June, respectively. In the case of rotavirus the incidence rate is higher in spring with a peak from March to April. The learned trend components capture the disease-specific baseline rate of infections, which remains stable throughout the years 2013 to 2016. While there is little uncertainty in the seasonal component, there is a high degree of uncertainty in the constant offset of the trend component. The effect of combining both contributions within the model’s exponential nonlinearity results in higher uncertainty around larger values.

**Fig 4.**
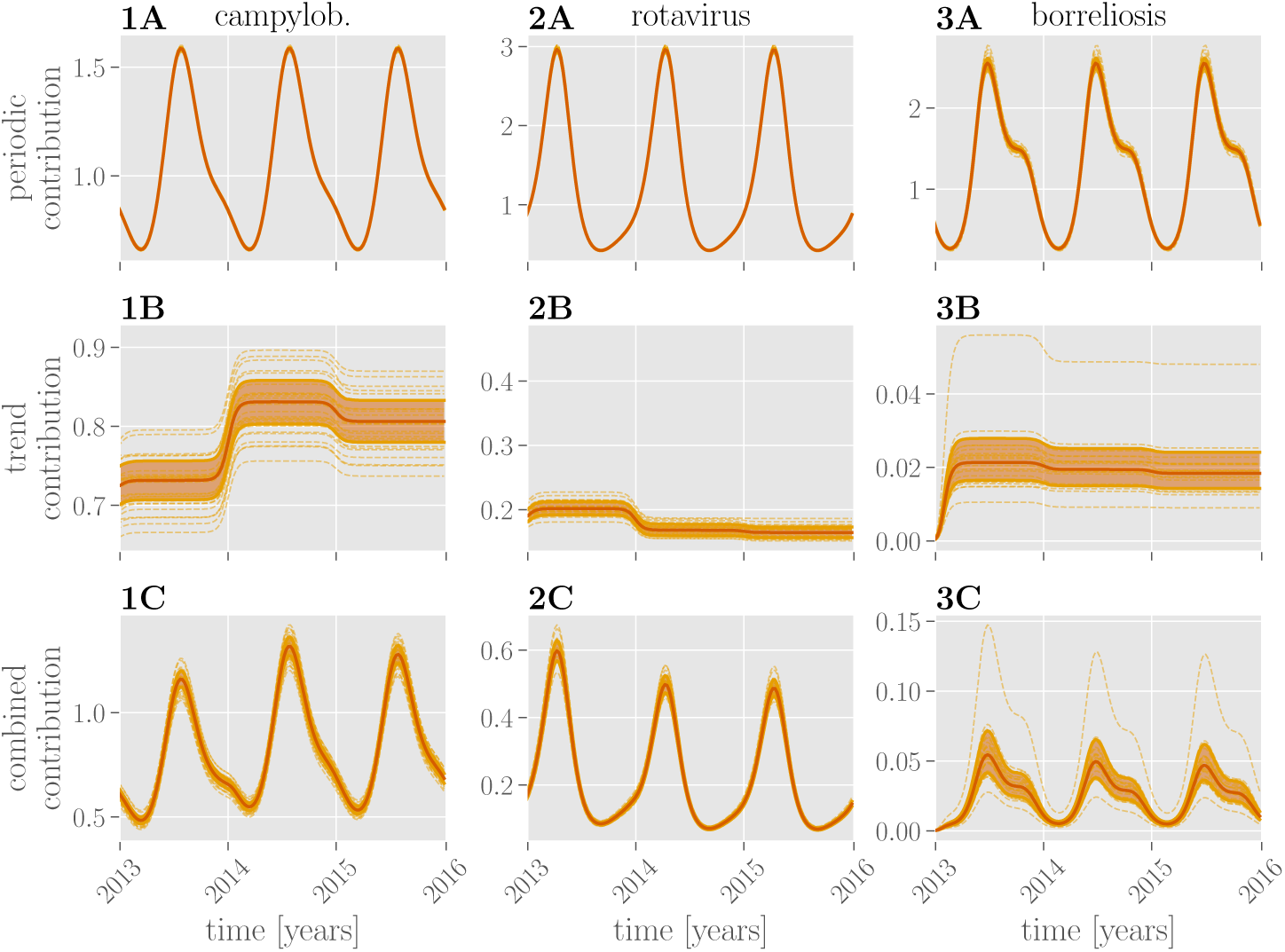
Learned temporal contributions. Periodic contributions over the course of three years (2013-2016) for all three diseases are shown in the **row A**, trend contributions in the **row B** and their combination in the **row C**. Red lines show the mean exponentiated linear combination of periodic or trend or both features through the respective parameters. Dashed lines show random samples thereof; the shaded region marks the 25%-75% quantile.

For campylobacteriosis and, to a lesser extent, rotavirus reported incidence rates are higher in regions formerly belonging to eastern Germany (see Fig. 5). The parameters inferred for demographic components (see Fig. 5) show the role that age stratification plays for susceptibility. For all three diseases, a larger share of children and adolescents (ages 5-20 years) in the general population is indicative of increased incidence rates. Additionally, working-age adults (ages 20-65 years) appear to increase the incidence rate of borreliosis. It should be noted that this does not necessarily imply an increased susceptibility of the respective groups themselves, but could instead be due to latent variables correlated with age stratification, such as economic or cultural differences. The pairwise joint distributions reveal strong (anti-)correlations of the coefficients associated with the demographic and political components. E.g. the coefficient associated with age group [20-65) is strongly correlated with the coefficient associated with the east/west component, which implies ambiguity in the optimal choice of parameters.

**Fig 5.**
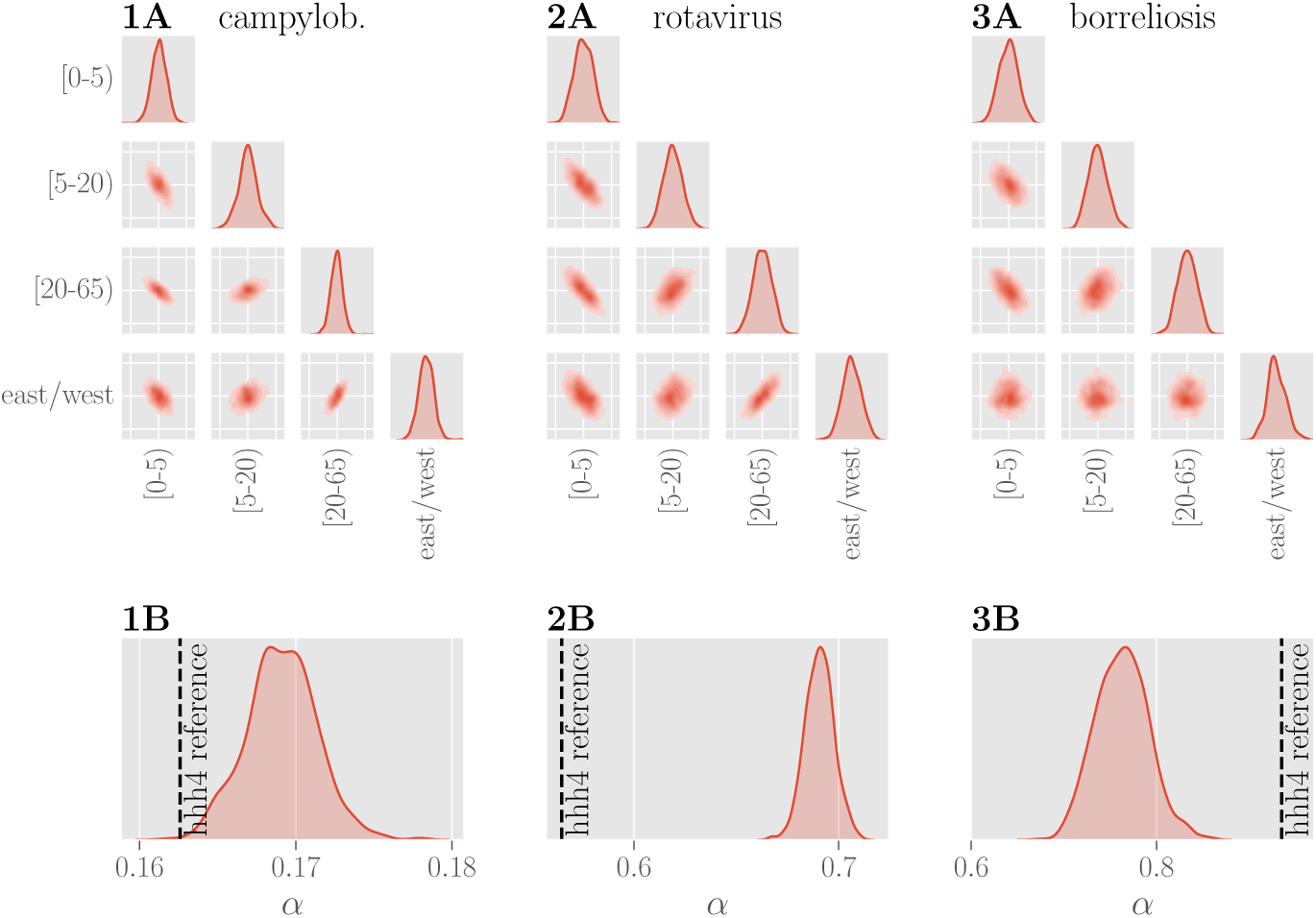
Learned weights for political and demographic components. Plots of the pairwise marginal distributions between inferred coefficients for three age groups and the east/west component for all three diseases are shown in **row A**. The marginal distribution of each coefficient shows a narrow unimodal peak, yet the pairwise distributions show that the individual features are clearly not independent. **Row B** shows the inferred posterior distributions of the overdispersion parameter *α* for three diseases. Values of *α* obtained using the *hhh4* reference model are indicated with a dashed black line. The inferred values for the dispersion parameter *α* are different, yet of similar magnitude, between the two models.

The posterior probability over the dispersion parameter *α* (see Fig.5) is tightly distributed around the respective disease specific means. With small values of *α*, the distribution of case counts for campylobacteriosis approaches a Poisson distribution, whereas the corresponding distributions for rotavirus and borreliosis are over-dispersed and deviate more from Poisson distributions.

### Predictive performance

The one-week-ahead predictions are shown in Fig.6, for two selected cities (Dortmund and Leipzig for campylobacteriosis and rotavirus, Nürnberg and München for borreliosis), together with the corresponding prediction from the reference *hhh4* model [18] fitted to the same data. A choropleth map of Germany (or the federal state of Bavaria in the case of borreliosis) shows the individual predictions for each county in one calendar week as an example. See also supplementary figures S4 Fig, S5 Fig and S6 Fig for predictions for 25 additional counties.

**Fig 6.**
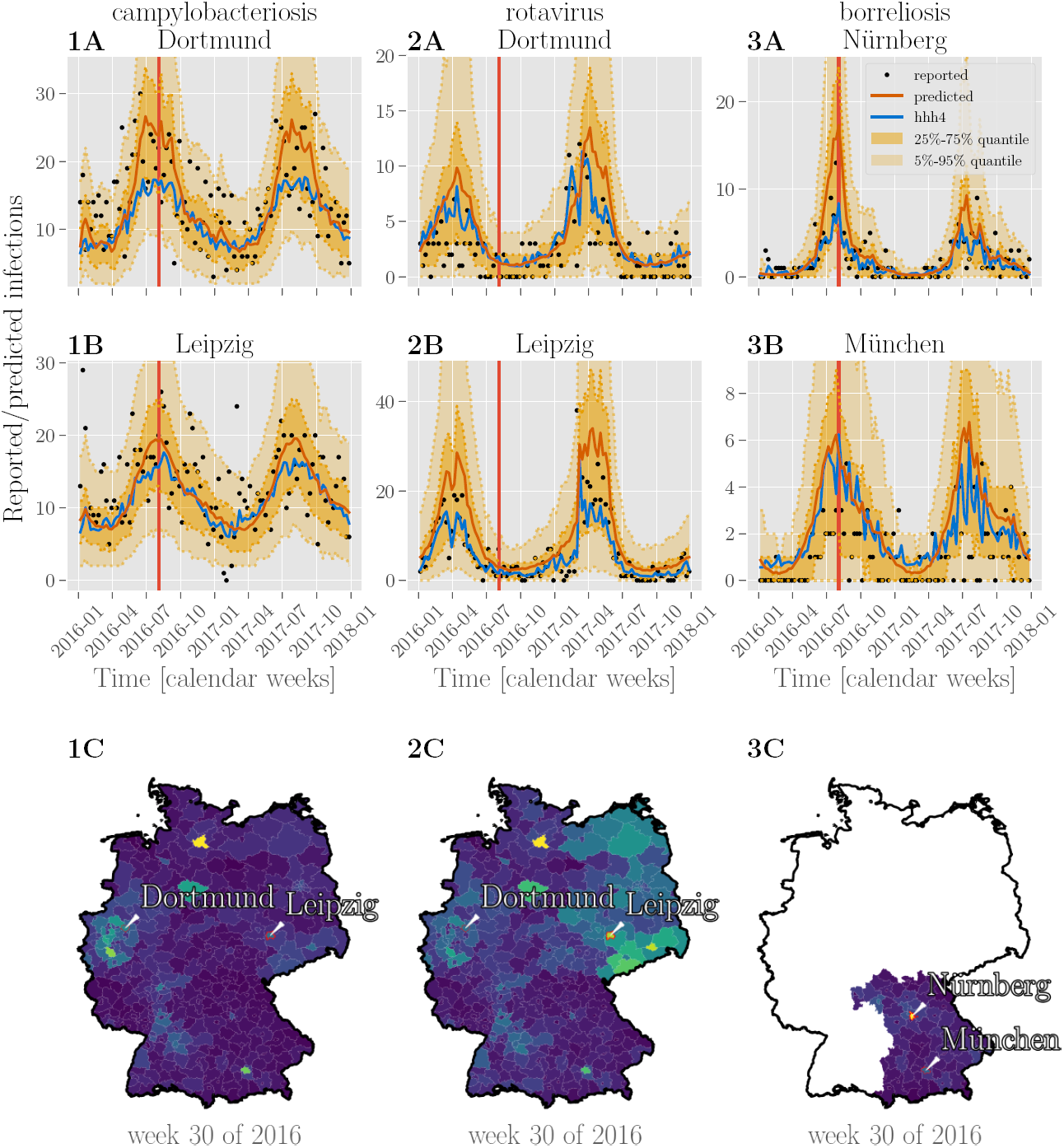
Predictions of case counts for various diseases by county. Reported infections (black dots), predictions of case counts by BSTIM (orange line) and the *hhh4* reference model (blue line) for campylobacteriosis **(column 1)**, rotavirus **(column 2)** and borreliosis **(column 3)** for two counties in Germany (for campylobacteriosis and rotavirus) or Bavaria (borreliosis), are shown in **rows A and B**. The shaded areas show the inner 25%-75% and 5%-95% percentile. **Row C** shows predictions of the respective disease for each county in Germany or the federal state of Bavaria in week 30 of 2016 (indicated by a vertical red line in rows A and B).

The BSTIM fits the mean of the underlying distribution of the data well. For rotavirus and borreliosis, it appears to overestimate the dispersion for the cities shown in Fig.6 as indicated by most data points falling within the inner 25%-75% quantile. This may be due to a too high dispersion parameter *α* (cf. Fig.5) or uncertainty about model parameters. It should be noted, however, that the optimal dispersion parameter itself varies from county to county, whereas our model infers only one single value for all counties together. The resulting predictions for all three diseases are smoother in time and space (cf. the chloropleth maps in Fig.6) than the predictions by the reference *hhh4* model. We attribute this to the smooth temporal basis functions and spatio-temporal interaction kernel of our model.

To quantitatively compare the performance of both models, we calculate the distributions of deviance and Dawid-Sebastiani score over all counties for BSTIM and the *hhh4* reference model as shown in Fig.7. Both measures show a very similar distribution of errors between both models for all three diseases, as it can be seen in table 2. Only for borreliosis, the *hhh4* model appears to be more sensitive to outliers.

**Fig 7.**
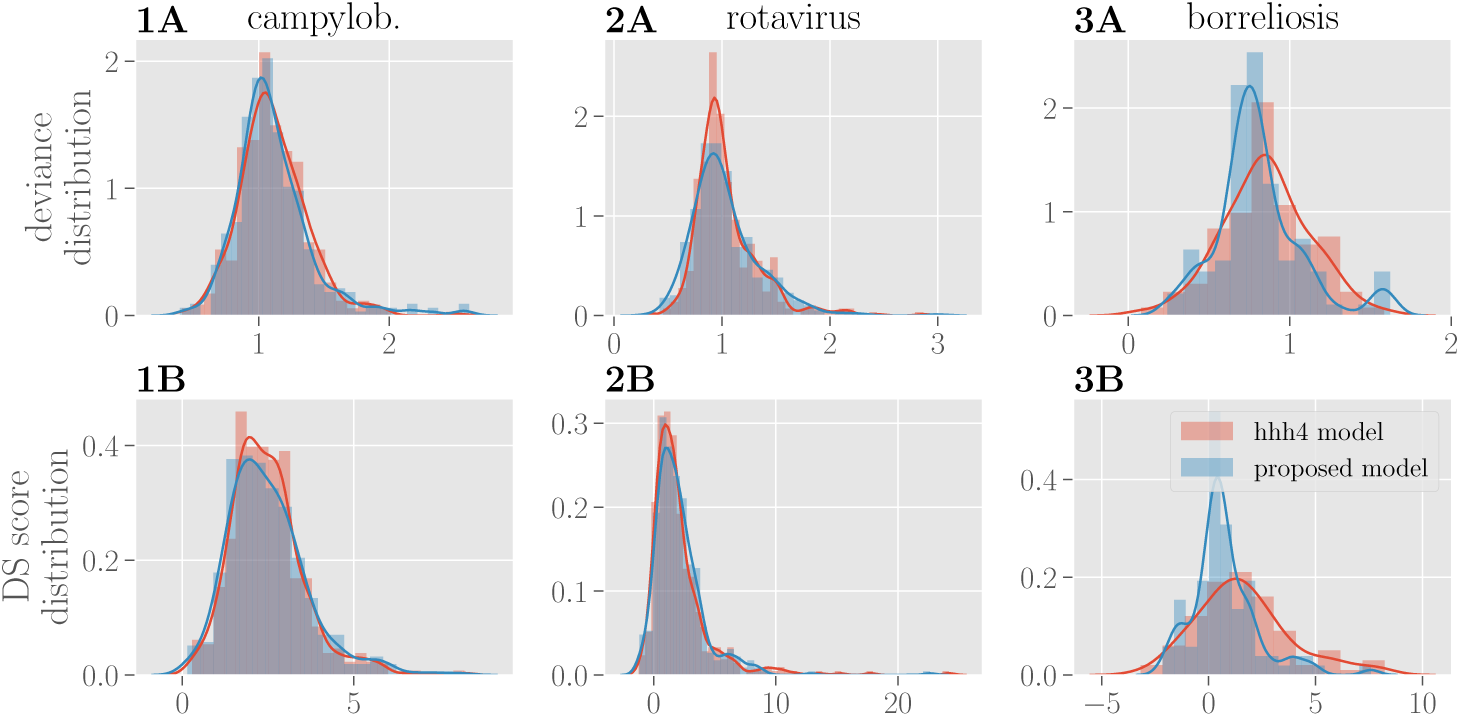
Evaluation of prediction performance. The distribution of deviance over counties is shown in **row A** for BSTIM (blue) and the reference *hhh4* model (red) for all three diseases. The corresponding distribution of Dawid-Sebastiani scores is shown in **row B**.

**Table 2.** Deviance and Dawid-Sebastiani score (mean ± standard deviation) for all three diseases and both BSTIM and the *hhh4* model.

| disease | score | BSTIM | *hhh4* |
| --- | --- | --- | --- |
| campylob. | deviance | $1.11 \pm 0.3$ | $1.11 \pm 0.26$ |
| | DS score | $2.49 \pm 1.18$ | $2.47 \pm 1.06$ |
| rotavirus | deviance | $1.03 \pm 0.32$ | $1.04 \pm 0.3$ |
| | DS score | $2.09 \pm 2.14$ | $2.1 \pm 2.54$ |
| borreliosis | deviance | $0.81 \pm 0.27$ | $0.85 \pm 0.27$ |
| | DS score | $0.74 \pm 1.56$ | $1.63 \pm 2.24$ |

### Benefits of probabilistic modeling for epidemiology

Probabilistic modeling relies on the specification of prior probability distributions over parameters [13]. In the context of epidemiology, this makes it possible to incorporate domain knowledge (e.g. we know that case counts tend to be overdispersed relative to Poisson distributions, but not to which degree for a specific disease) as well as modeling assumptions (e.g. we constrain interaction effects to be nonnegative). This is particularly relevant for diseases with limited available data (e.g. those not routinely recorded through surveillance), where appropriately chosen priors are required to prevent overfitting. The framework can easily be extended to include additional features or latent variables. For example, we introduce precise locations and times of individual cases as latent variables, given only the counties and calendar weeks in which they occurred.

Probabilistic models as discussed here provide samples of the posterior distribution of parameters as well as model predictions. This allows for analysis that is not possible with point estimation techniques such as maximum likelihood estimation. In epidemiology, datasets can be small, noisy or collected with low spatial or temporal resolution. This can lead to ambiguity, where the observations could be equally well attributed to different features and thus different model parameterizations are plausible. While maximum likelihood estimation in such a situation selects only the single most likely model, Bayesian modeling captures the full distribution over possible parameters and predictions, and thus preserves information about the uncertainty associated with the parameters of the model itself. Analyzing the parameter distribution can thus help identify redundant or uninformative features.

Samples from the inferred parameter distributions are afterwards used to derive samples of predicted future cases. The resulting predictions thus incorporate both noise assumptions about the data as well as model uncertainty. This can be relevant for determining confidence intervals, in particular in situations where model uncertainty is large. The samples of the predictive distribution can in turn be used for additional processing, or if predictions in the form of point estimates are desired, they can be summarized by the posterior mean.

### Possible extensions

Due to the flexibility of the probabilistic modeling and sampling approach, additional variables can be easily included and their influence analyzed (e.g. weather data, geographical features like forests, mountains and water bodies, the location and size of hospitals, vaccination rates, migration statistics, socioeconomic features, population densities, self-reported infections on social media [19], work, school and national holidays, weekends and large public events). For features where precise values are not known, probability distributions could be specified and included in the probabilistic model, which could improve the model’s estimate of uncertainty. Whereas spatio-temporal interaction effects are here modeled as a function of geographical proximity, the kernel’s composite basis functions make it possible to use alternative spatial distance measures such as derived from transportation networks for people or goods [20]. Since the precise locations and times of individual infections are not publicly known, we simply assume a geographically and temporally uniform distribution of cases within the given county and calendar week. The conditional probability distributions could be refined by incorporating additional information (e.g. weekends and population density maps). However, precise information on place and time of infection are available to local health agencies: The model presented here could readily be implemented there and use exact space and time data.

## Conclusion

In this paper, a probabilistic model is proposed for predicting case counts of epidemic diseases. It takes into account a history of reported cases in a spatially extended region and employs MCMC sampling techniques to derive posterior parameter distributions, which in turn are incorporated in predicted probability distributions of future infection counts across time and space.

For all three tested diseases (campylobacteriosis, rotavirus and borreliosis) the same model, using interaction effects, temporal, political and demographic information, performs well and produces smooth predictions in time and space.

A comparison with the standard *hhh4* model, which uses maximum likelihood estimation instead of Bayesian inference, shows comparable performance. At the expense of higher computational costs than the point estimate used in *hhh4*, the sampling approach employed here provides information about the full posterior distribution of parameters and predictions. The posterior parameter distribution provides information about the relevance of the corresponding features for the inferred model, and helps in identifying redundant features or violated model assumptions. The inferred features of our model are interpretable and their individual contribution to the model prediction can be analyzed: spatio-temporal interactions reveal information about the dynamic spread of the disease, temporal features capture seasonal fluctuations and long-term trends, and the assigned weights indicate relevance of additional features. The posterior predictive distribution also accounts for the uncertainty about parameters, due to simplifying model assumptions or a lack of data, rather than just the variability inherent in the data itself. This additional information is valuable for public-health policy-making, where accurate quantification of uncertainty is critical.

## Supporting information

**S1 Fig.**
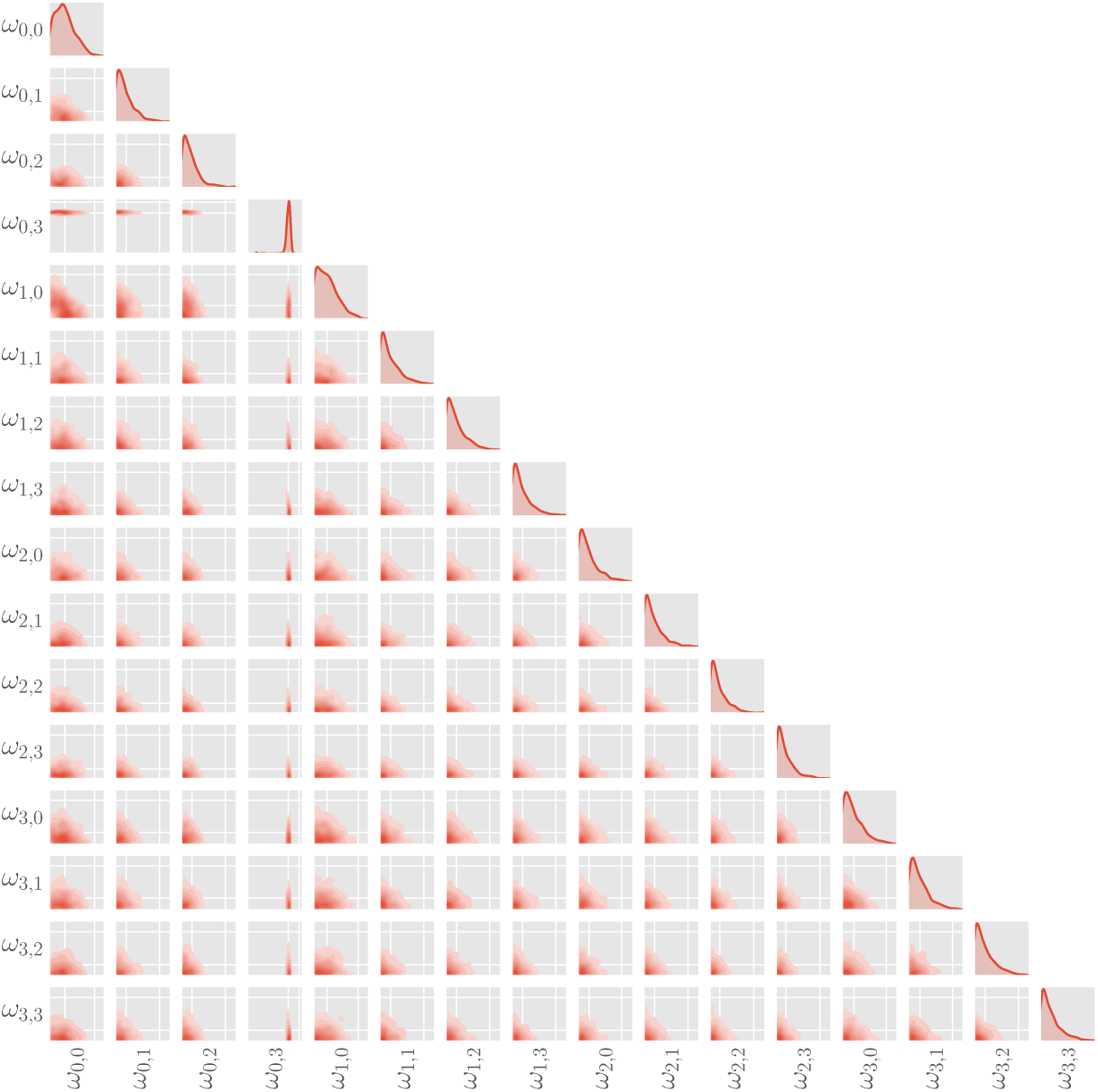
Pairwise correlation plot of weights for interaction components inferred for campylobacteriosis. Each sub-diagonal plot shows the (marginal) joint distribution and regression line for two coefficients of the interaction kernel inferred from training data for campylobacteriosis. The plots on the diagonal show the respective univariate marginal distributions of each coefficient. Note that since the coefficients are constrained to positive values, only the first quadrant is shown.

**S2 Fig.**
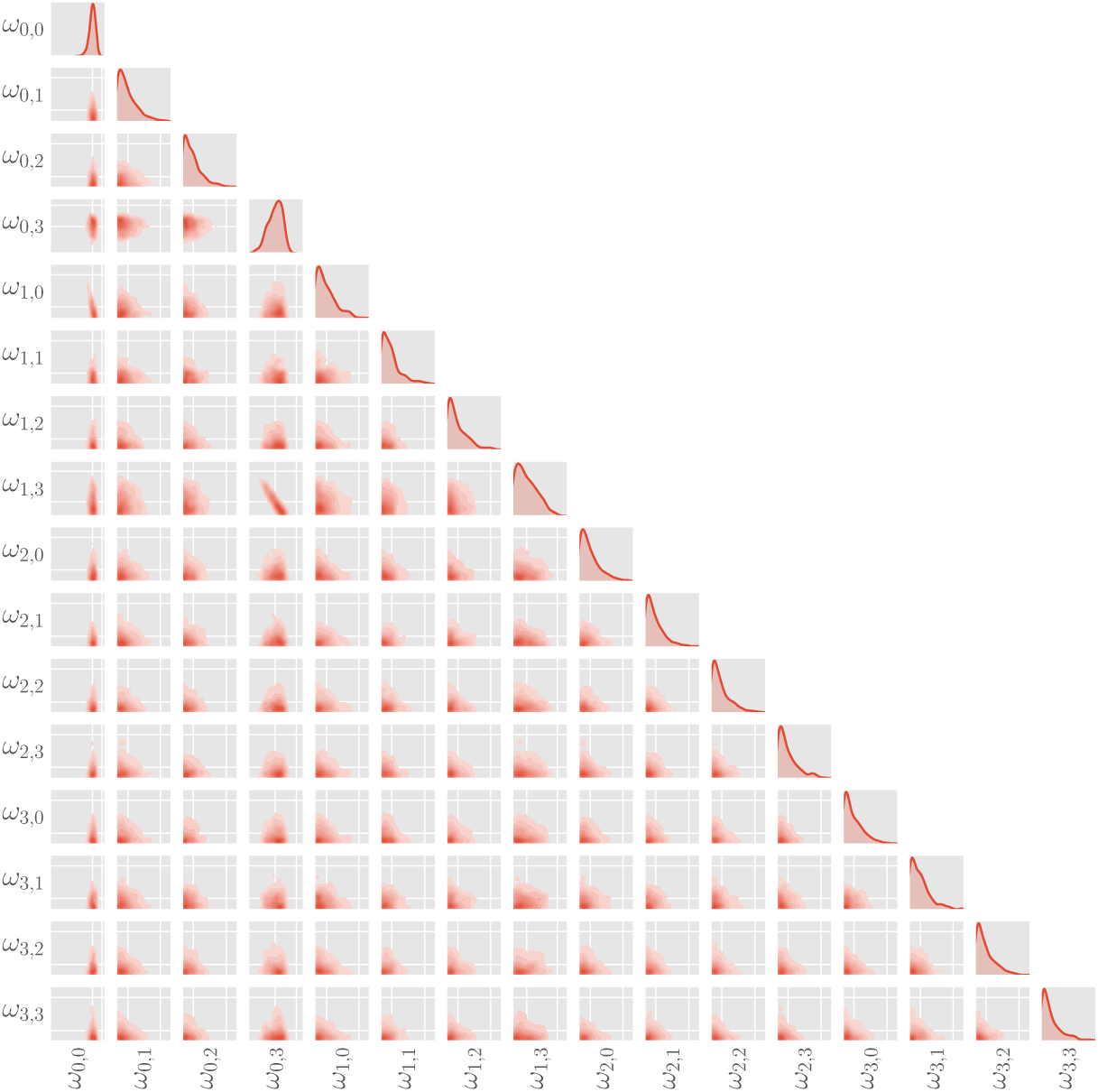
Correlation plot of weights for interaction components inferred for rotavirus. Each sub-diagonal plot shows the (marginal) joint distribution and regression line for two coefficients of the interaction kernel inferred from training data for rotavirus. The plots on the diagonal show the respective univariate marginal distributions of each coefficient. Note that since the coefficients are constrained to positive values, only the first quadrant is shown.

**S3 Fig.**
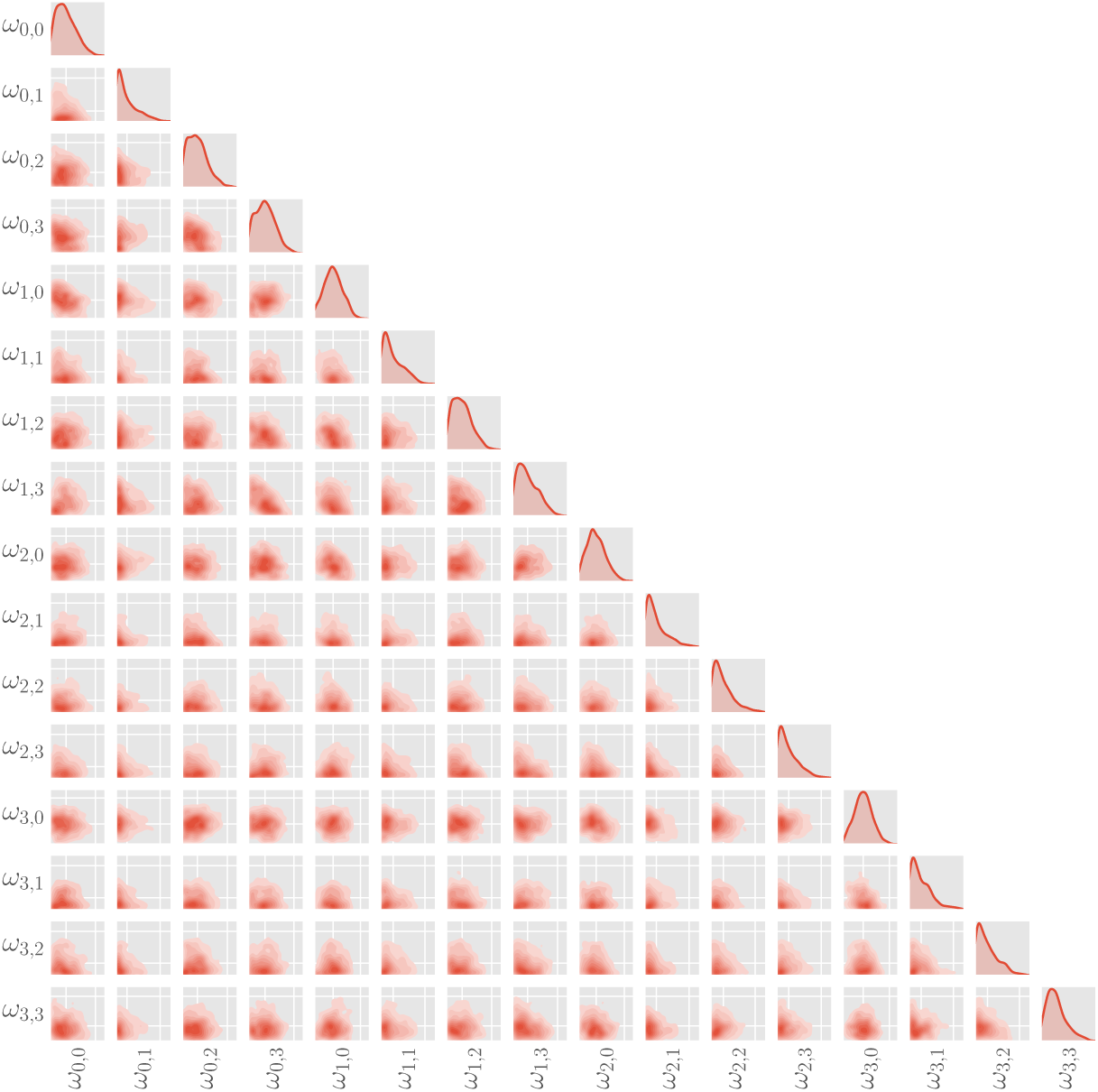
Correlation plot of weights for interaction components inferred for borreliosis. Each sub-diagonal plot shows the (marginal) joint distribution and regression line for two coefficients of the interaction kernel inferred from training data for borreliosis. The plots on the diagonal show the respective univariate marginal distributions of each coefficient. Note that since the coefficients are constrained to positive values, only the first quadrant is shown.

**S4 Fig.**
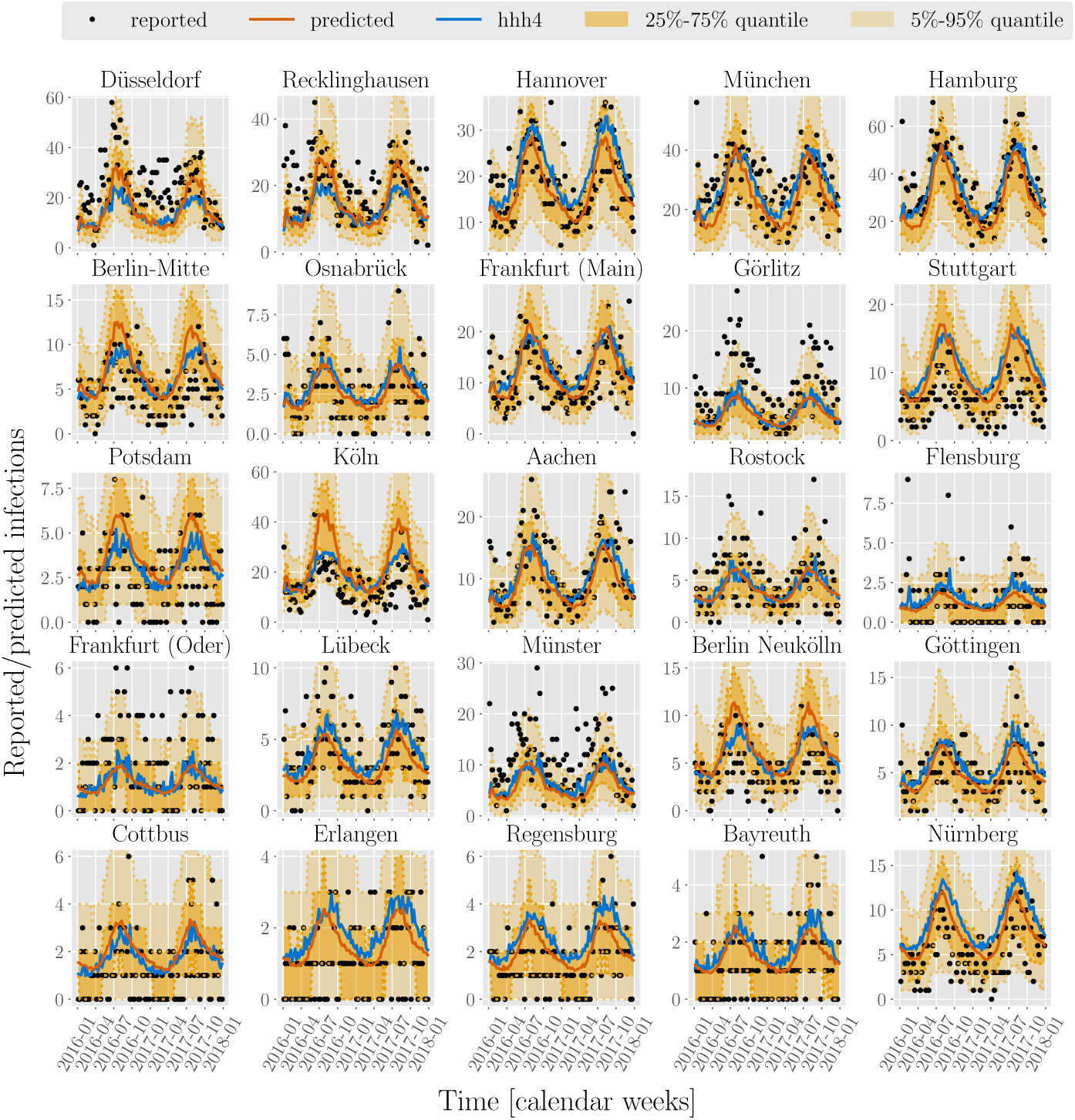
Predictions of case counts for campylobacteriosis for various counties across Germany. Reported infections (black dots), predictions of case counts by BSTIM (orange line) and the *hhh4* reference model (blue line) for campylobacteriosis for 25 counties in Germany. The shaded areas show the inner 25%-75% and 5%-95% percentile.

**S5 Fig.**
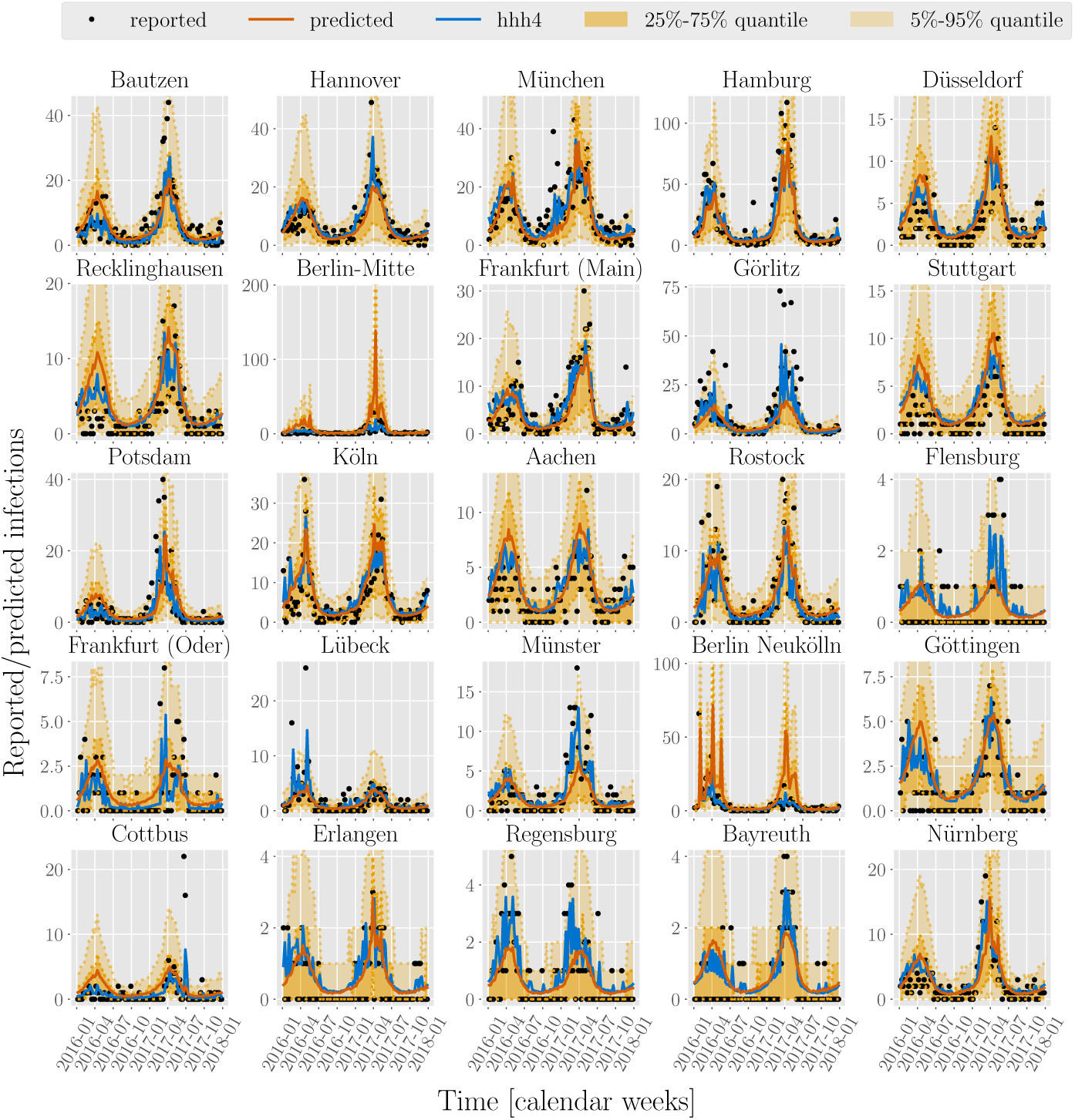
Predictions of case counts for rotavirus for various counties across Germany. Reported infections (black dots), predictions of case counts by BSTIM (orange line) and the *hhh4* reference model (blue line) for rotavirus for 25 counties in Germany. The shaded areas show the inner 25%-75% and 5%-95% percentile.

**S6 Fig.**
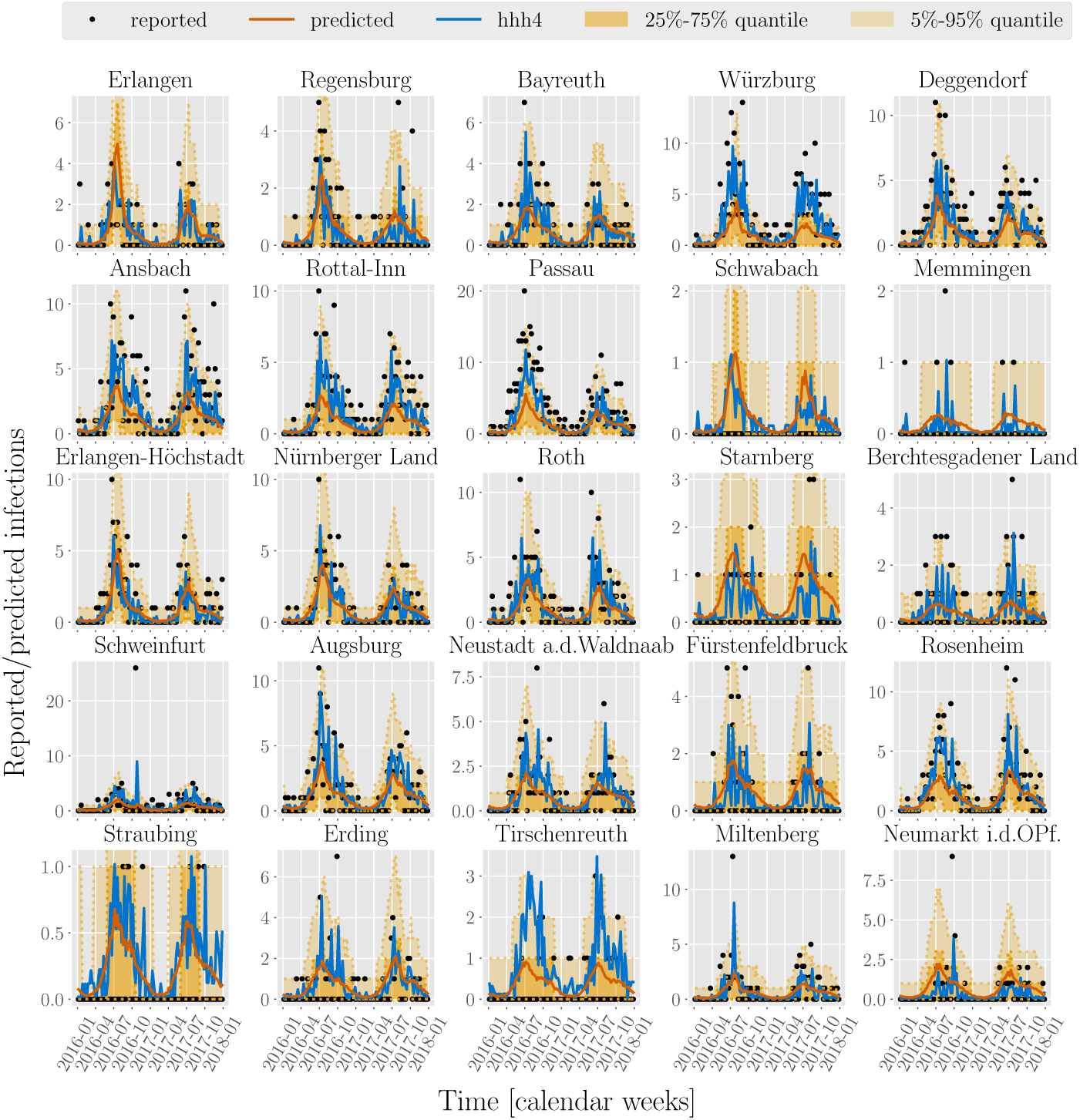
Predictions of case counts for borreliosis for various counties across Bavaria. Reported infections (black dots), predictions of case counts by BSTIM (orange line) and the *hhh4* reference model (blue line) for borreliosis for 25 counties in Bavaria. The shaded areas show the inner 25%-75% and 5%-95% percentile.

## Footnotes

1 We use the term “county” to generally refer to rural counties (*Landkreise*) and cities (*kreisfreie Städte*) as well as the twelve districts of Berlin (*Bezirke*).

2 The age group of 65 years and above accounts for the remaining share of the population and thus is a redundant variable with respect to the other three age groups and the total population.

